# *In vivo* calcium imaging of CA3 pyramidal neuron populations in adult mouse hippocampus

**DOI:** 10.1101/2021.01.21.427642

**Authors:** Gwendolin Schoenfeld, Stefano Carta, Peter Rupprecht, Aslı Ayaz, Fritjof Helmchen

## Abstract

Neuronal population activity in the hippocampal CA3 subfield is implicated in cognitive brain functions such as memory processing and spatial navigation. However, because of its deep location in the brain, the CA3 area has been difficult to target with modern calcium imaging approaches. Here, we achieved chronic two-photon calcium imaging of CA3 pyramidal neurons with the red fluorescent calcium indicator R-CaMP1.07 in anesthetized and awake mice. We characterize CA3 neuronal activity at both the single-cell and population level and assess its stability across multiple imaging days. During both anesthesia and wakefulness, nearly all CA3 pyramidal neurons displayed calcium transients. Most of the calcium transients were consistent with a high incidence of bursts of action potentials, based on calibration measurements using simultaneous juxtacellular recordings and calcium imaging. In awake mice, we found state-dependent differences with striking large and prolonged calcium transients during locomotion. We estimate that trains of >30 action potentials over 3 s underlie these salient events. Their abundance in particular subsets of neurons was relatively stable across days. At the population level, we found that coactivity within the CA3 network was above chance level and that co-active neuron pairs maintained their correlated activity over days. Our results corroborate the notion of state-dependent spatiotemporal activity patterns in the recurrent network of CA3 and demonstrate that at least some features of population activity, namely coactivity of cell pairs and likelihood to engage in prolonged high activity, are maintained over days.

## INTRODUCTION

Neuronal populations in the hippocampal CA3 subfield are part of the mammalian brain circuit that is essential for spatial navigation, memory formation, and cognition (Kesner, 2007; Hartley et al., 2013; Rolls, 2016; Rebola et al., 2017; Hainmueller and Bartos, 2018). CA3 pyramidal neurons are special in forming an auto-associative recurrent network enabling memory encoding and pattern completion (Rolls, 2007; Kesner and Rolls, 2014; Guzman et al., 2016; Knierim and Neunuebel, 2016). The functional properties of CA3 pyramidal neurons have been characterized largely with electrophysiology, using extracellular recordings (Fox and Ranck, 1975; Csicsvari et al., 2000; Henze et al., 2002; Leutgeb et al., 2004; Frerking et al., 2005; Mizuseki et al., 2012, Oliva et al., 2016), in vivo intra- and juxtacellular recordings (Epsztein et al., 2011; Kowalski et al., 2016; Zucca et al., 2017; Diamantaki et al., 2018; Hunt et al., 2018; Malezieux et al., 2020), and whole-cell recordings in brain slices (Jonas et al., 1993; Hemond et al., 2008; Hunt et al., 2018; Raus Balind et al., 2019). Pyramidal neurons in CA3 show properties distinct from CA1 (Mizuseki et al., 2012; Oliva et al., 2016) but display heterogeneity within their population (Cembrowski and Spruston, 2019; Ding et al., 2020; Hunt et al., 2018). For CA3 pyramidal neurons, mean firing rates typically range from 0.3 to 5 Hz in vivo (Henze et al., 2002; Wittner and Miles, 2007; Mizuseki et al., 2012; Kowalski et al., 2016; Oliva et al., 2016; Ding et al., 2020), lower than for CA1 pyramidal neurons but higher when compared to dentate gyrus (DG) granule cells. As a prominent feature, hippocampal pyramidal neurons, especially in CA3, exhibit bursts of action potentials (APs) with inter-spike intervals <6 ms (Fox and Ranck, 1975; Frerking et al., 2005; Mizuseki et al., 2012; Kowalski et al., 2016; Oliva et al., 2016; Raus Balind et al., 2019). These complex spike bursts involve regenerative dendritic mechanisms and have been implicated in activity-dependent plasticity (Lee et al., 2012; Grienberger et al., 2014; Bittner et al., 2017, 2015; Diamantaki et al., 2018; Raus Balind et al., 2019). They are also associated with network synchronization events in CA3 (Miles and Wong, 1983; Menendez De La Prida et al., 2006; Wittner and Miles, 2007; Marissal et al., 2012), especially sharp-wave ripples (Buzsáki, 1986; Csicsvari et al., 2000; Harris et al., 2003; Hunt et al., 2018).

Despite these advances in electrophysiological studies, our understanding of CA3 network dynamics and its computational roles remains limited. Optophysiology offers promising complementary approaches, especially in terms of longitudinal imaging of the same neuronal population. However, due to the difficulties in accessing deeper brain regions, hippocampal imaging studies have lagged behind similar studies in neocortex. Only during the last decade, in vivo calcium imaging in hippocampus became possible, typically by removing the overlying cortical tissue and using either two-photon microscopy in head-fixed animals (Dombeck et al., 2010; Grienberger et al., 2014; Hainmueller and Bartos, 2018; Kinsky et al., 2018) or mini-endoscopes in freely-moving mice (Ziv et al., 2013; Rubin et al., 2015; Gonzalez et al., 2019; Stefanini et al., 2020). While initial studies mainly targeted CA1 as the most accessible region, only at a later stage chronic and functional imaging was also established in the DG (Pilz et al., 2016, 2018; Danielson et al., 2017; Hainmueller and Bartos, 2018; Stefanini et al., 2020). In our own previous study (Pilz et al., 2016), by applying GCaMP6 and specifically R-CaMP1.07, a red calcium indicator that facilitates deep imaging (Ohkura et al., 2012; Bethge et al., 2017), we confirmed sparse activity of DG granule cells and described its variation across behavioral states. Functional imaging in CA3 is as challenging as in DG and therefore has been achieved in only few studies until today (Rajasethupathy et al., 2015; Hainmueller and Bartos, 2018; Rashid et al., 2020). As an emerging field, CA3 imaging provides new opportunities to address key questions about cellular and circuit mechanisms of neural coding and plasticity in this region.

Here, we establish in vivo calcium imaging of CA3 pyramidal neurons using an approach similar to our previous DG study (Pilz et al., 2016). We characterize basic features of CA3 calcium transients and calibrate them in terms of underlying APs using simultaneous juxtacellular recordings. We find heterogeneous CA3 activity patterns across behavioral states and discover particularly prominent prolonged calcium transients that occur in neuronal subsets during running. Moreover, our longitudinal imaging results indicate that CA3 population activity at least partially remains stable across days, particularly with respect to the co-activity of neurons within sub-ensembles.

## RESULTS

### In vivo two-photon calcium imaging of CA3 pyramidal neurons

We established in vivo two-photon imaging of CA3 neuronal population activity through a chronically implanted window after removal of cortical tissue overlying the hippocampus (Fig. 1A, B). To induce expression of a genetically encoded calcium indicator specifically in CA3 pyramidal neurons, we injected Girk4-Cre transgenic mice a virus driving Cre-dependent expression of the red-shifted calcium indicator R-CaMP1.07 (Ohkura et al., 2012; Bethge et al., 2017) (Fig. 1B). Following chronic window preparation and habituation of the mouse to head-fixation, we performed calcium imaging of R-CaMP1.07-expressing CA3 pyramidal neurons in several sessions of around 30-min duration, spread over consecutive days and under either anesthetized or awake condition (Fig. 1C). During awake recordings, mice were free to run or rest on a ladder wheel placed under the two-photon microscope. We continually measured running speed and used a threshold to distinguish behavioral states by defining ‘run’ and ‘rest’ periods.

**Figure 1.**
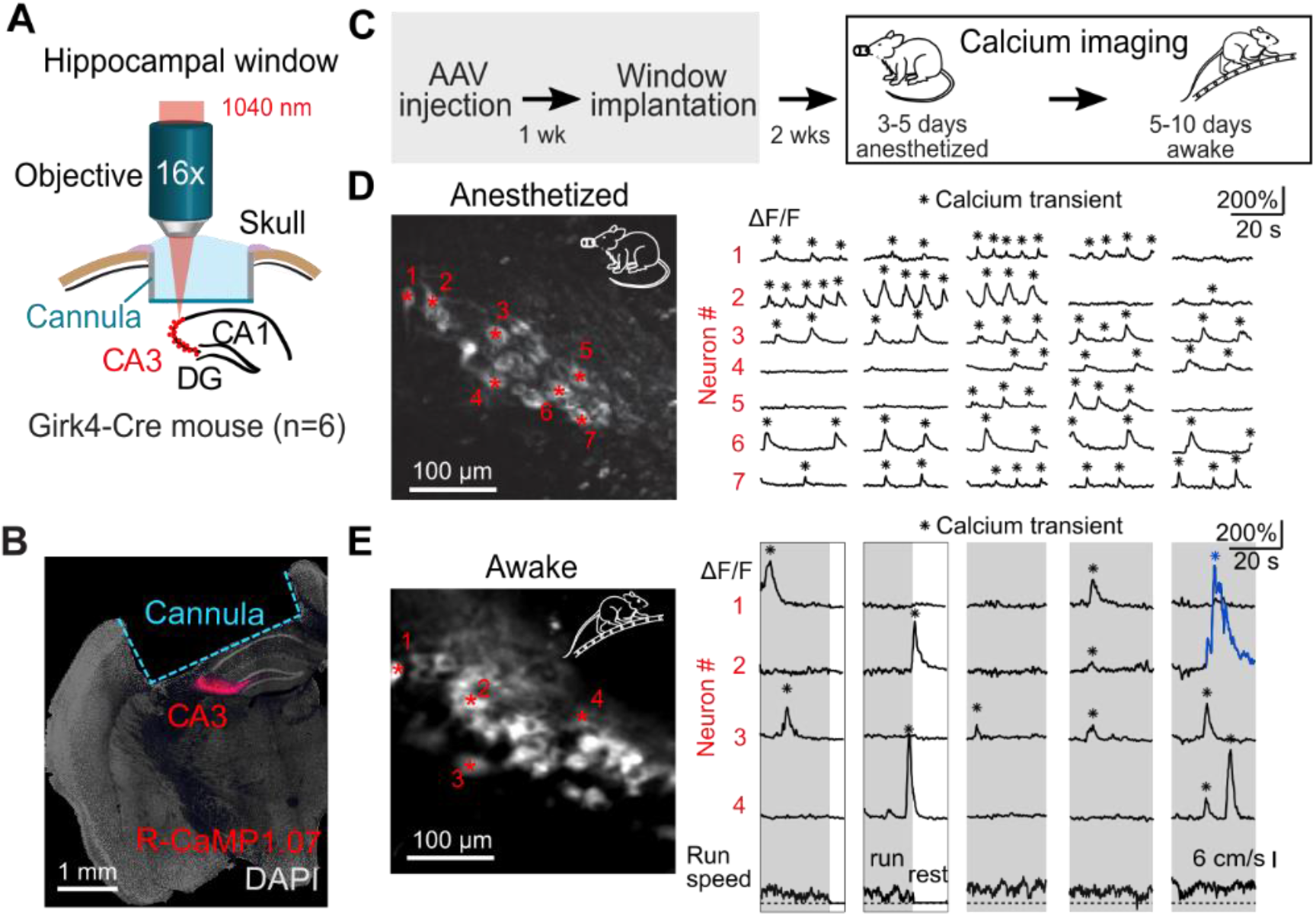
*In vivo* two-photon imaging of CA3 pyramidal neuron populations during wakefulness and anesthesia. ***A***, Schematic illustration of experimental setup showing the chronic window implant above the corpus callosum and area CA1 of the intact hippocampus for R-CaMP1.07 calcium imaging in CA3. **B**, Histological coronal cross-section of the fixed brain after *in vivo* imaging sessions showing R-CaMP1.07 labelled CA3 pyramidal neurons and the hippocampal window implant. ***C***, Experimental time line. After recovery from the surgery, three consecutive days of anesthetized imaging were carried out. This was followed by six to ten days of awake imaging including at least three consecutive imaging days. ***D***, Example field of view of an anesthetized two-photon imaging session. Example neurons are labelled by red asterisks and their respective ΔF/F traces are displayed on the right. Detected calcium transient peaks are labelled with black asterisks. ***E***, Example field of view and ΔF/F traces of neurons recorded during wakefulness. An example large calcium transient is labelled in blue. Run speed of the animal is shown below.

Nearly all neurons exhibited calcium transients indicative of neuronal spiking activity in both anesthetized and awake condition (Fig. 1D, E). For every mouse (n = 6) more than 90% of cells showed at least one detectable calcium transient on the first imaging day with isoflurane-anesthesia (92%, 142 cells, 6 FOVs) as well as on the first awake imaging day (96%, 355 cells, 10 FOVs). Calcium transients occurred rather regularly in individual neurons in anesthetized mice. In contrast, amplitudes and durations of calcium transients were more heterogeneous in awake mice, including a substantial fraction of large and prolonged events (an example is colored in Fig. 1E). On average, calcium transients were smaller and of shorter duration during anesthesia compared to wakefulness (ΔF/F peak amplitude 45.0 ± 26.3% vs. 89.5 ± 65.0%; full-width-half-maximum [FWHM] 1.8 ± 2.3 s vs. 2.3 ± 2.1 s; mean ± s.d., 2934 transients in 382 neurons for anesthetized and 2806 transients in 388 neurons for awake condition; p < 0.001 for both comparisons; two-sided Wilcoxon rank sum test). These results indicate that CA3 pyramidal neurons show distinct patterns of neuronal activity in anesthetized compared to awake condition.

### Juxtacellular recordings of R-CaMP1.07 expressing CA3 pyramidal neurons

To relate R-CaMP1.07 calcium transients to AP patterns we performed acute experiments in anesthetized mice, obtaining simultaneous juxtacellular recordings and functional calcium imaging data from R-CaMP1.07-expressing CA3 pyramidal neurons (Fig. 2A, B). We extracted spike times using simple thresholding and temporally aligned calcium transients to the voltage recordings. Juxtacellular recordings revealed APs in variable numbers, often occurring in high-frequency bursts (Fig. 2C). The amplitude of consecutive spikes within a burst decreased over 4-6 APs, until no more spikes could be detected. For longer bursts, the AP amplitude often partially recovered after this initial decrease (Fig. 2C; Extended Data Fig. 2-1A, B). Burstiness was apparent in the distribution of inter-spike intervals (ISI) with a mean of 0.34 s, reflecting the inverse spike rate, and a median of 5.5 ms, indicating that two subsequent spikes within a burst typically were separated by <6 ms (Extended Data Fig. 2-1C).

**Figure 2.**
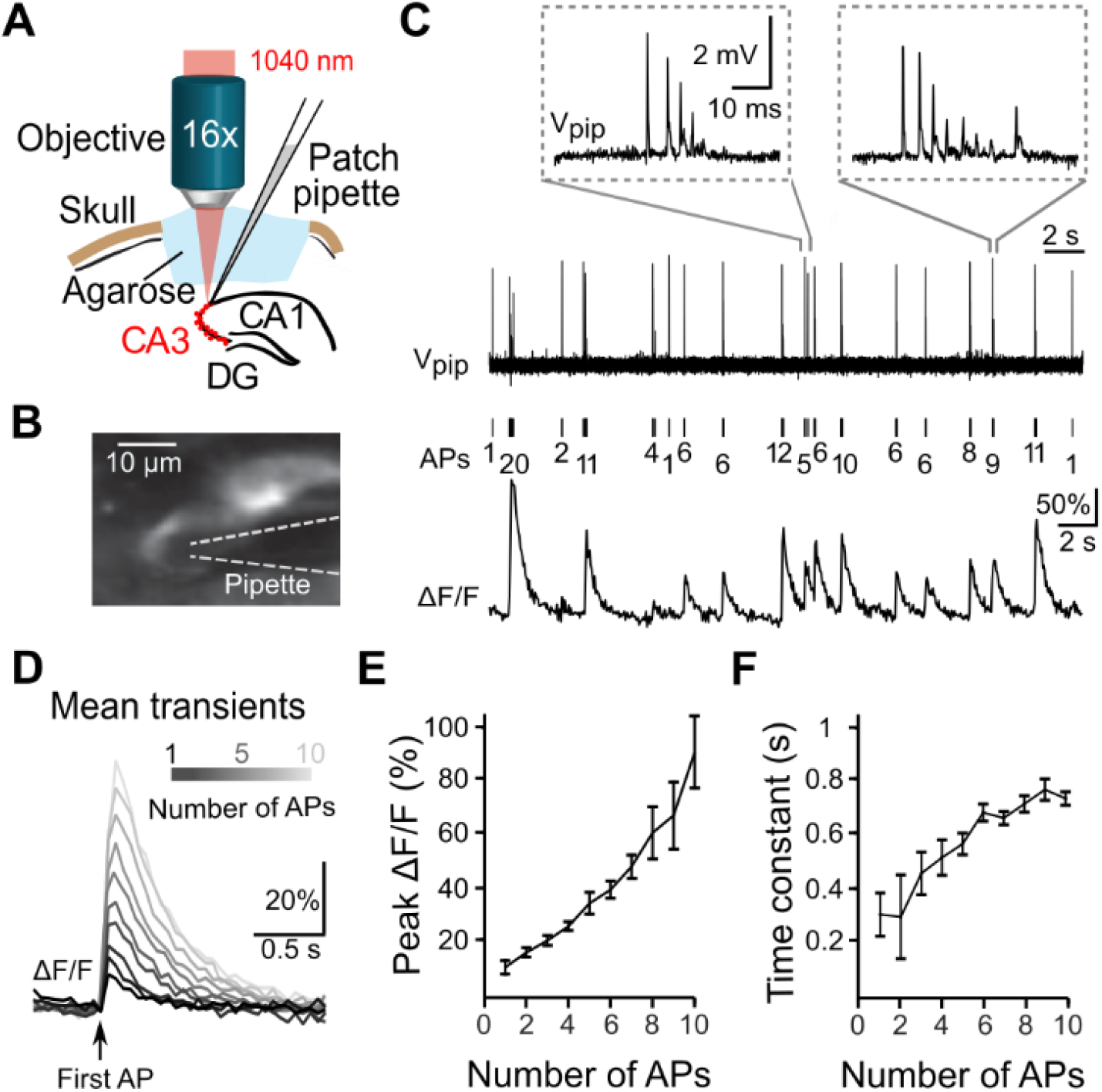
Calibration of *in vivo* spiking activity of CA3 pyramidal neurons using simultaneous two-photon calcium imaging and juxtacellular voltage recordings in acute experiments. ***A***, Schematic of experimental setup. ***B***, Maximum-intensity projection of two-photon images of a R-CaMP1.07-labelled CA3 pyramidal neurons together with the recording pipette in juxtacellular configuration. ***C***, Example traces of simultaneously recorded calcium transients (ΔF/F) and spontaneous APs (V_pip_) during isoflurane-anesthesia. The number of APs per burst is indicated below. Inserts show two example burst events magnified. ***D***, Average calcium transients caused by 1 to 10 APs aligned to the occurrence of the first AP. ***E***, Relationship between peak ΔF/F changes and number of APs (mean ± SEM). ***F***, Relationship between the calcium transient decay time constant and the number of APs (exponential fits; error bars indicate 90% confidence interval).

AP patterns in individual neurons correlated with the measured calcium transients (Fig. 2C). A spontaneous single AP-evoked ΔF/F transient on average had a peak amplitude of 11 ± 3% (n = 47 events, 4 neurons, 3 mice). With increasing number of APs, the ΔF/F amplitude of the corresponding calcium transients increased, following an approximately linear relationship up to 10 APs (Fig. 2D-E). The time constant of the evoked transient, as measured by an exponential fit, was around 0.3 s for single APs and remained <0.8 s for larger numbers of APs (Fig. 2F). These ground-truth data are an important calibration resource that helps to interpret R-CaMP1.07 imaging data in CA3 neuronal populations more quantitatively.

### Large and prolonged calcium transients during wakefulness and locomotion

Taking advantage of this ground-truth calibration we trained a supervised spike inference algorithm based on a deep neuronal network (Rupprecht et al., 2020) to temporally deconvolve ΔF/F transients and infer instantaneous spike rates (SRs) (Methods). Deconvolution uncovered that during wakefulness, in contrast to anesthesia, calcium transients often were prolonged, indicating extended periods of spiking, sometimes over seconds (Fig. 3A). For quantification, we computed the mean ΔF/F value in a 3-s time window around the calcium transient peak (1 s before until 2 s after the peak), reflecting the integral cellular activity causing the calcium transient. On average, the mean ΔF/F level was significantly higher during wakefulness compared to anesthesia (Fig. 3B; 23 ± 24% vs. 11 ± 10%, mean ± s.d., p<0.001, two-sided Wilcoxon rank sum test), in qualitative agreement with recent findings in CA1 (Yang et al., 2020). The distribution of mean ΔF/F values for the awake condition showed a pronounced tail of large events, with a substantial fraction reaching >100% mean ΔF/F. To account for these special events, we defined ‘awake large events’ as those calcium transients with mean ΔF/F values larger than most anesthetized events (>95th percentile; Fig. 3B). According to this definition, 33.7% of all awake events were classified as large events. Overall, we classified our recorded calcium transients in ‘anesthetized’ (n = 2934), awake ‘small’ (n = 1859) and awake ‘large’ (n = 947) events. We did not further divide calcium transients that were measured during anesthesia into small and large transients.

**Figure 3.**
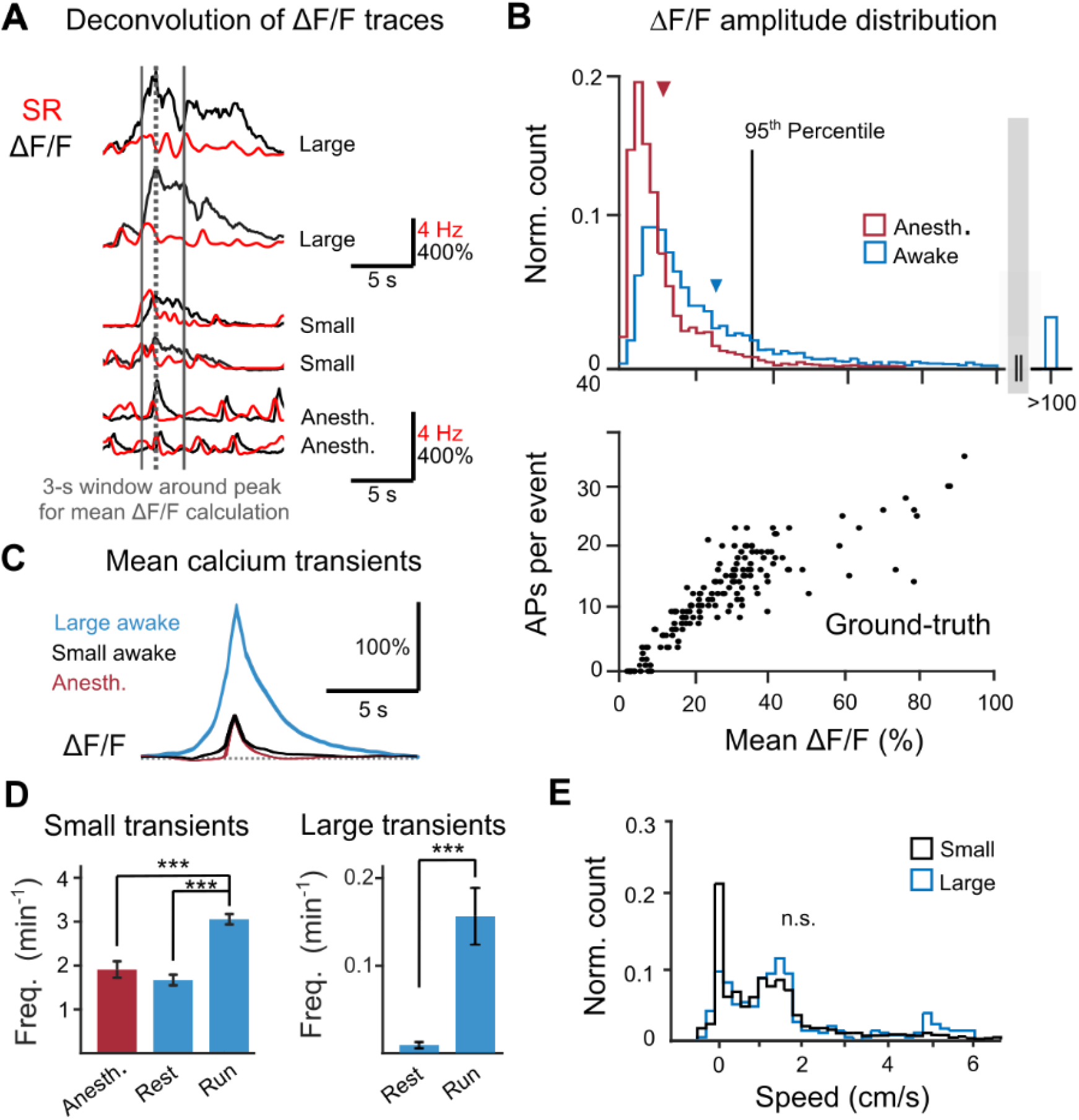
Large and prolonged calcium events in CA3 pyramidal neurons during wakefulness. ***A***, Example ΔF/F traces (black) with corresponding deconvolved instantaneous firing rates (red). ***B***, Distribution of mean ΔF/F level in a 3-s time window around the transient peak (see A) under anesthesia (red) and during wakefulness (blue). Wakeful transients that exceeded the 95^th^ percentile (37% ΔF/F) of anesthetized mean ΔF/F levels were classified as large events. Arrows indicate the respective mean of each distribution (awake: 23 ± 24%, anesthetized: 11 ± 10%, mean ± s.d., p<0.001, two-sided Wilcoxon rank sum test). The lower panel, as a calibration look-up table for the upper panel, shows the relation of mean ΔF/F level to the number of action potentials extracted from the ground-truth data set. ***C***, Average shape of calcium transients classified as anesthetized events, awake small and awake large events (mean ± SEM). ***D***, *Left:* Average frequency of events occurring under anesthesia (n=2934 events), of small wakeful (n=1859 events) and large wakeful events (n=947 events) (mean ± SEM). *Right:* Average frequency of small and large events under anesthesia and during wakeful resting and wakeful locomotion (mean ± SEM). ***p <0.001, one-way ANOVA (small events: p_anesth./run_ = 1.7×10^−4^, p_rest/run_ = 4.5×10^−16^; large events: p_rest/run_ = 6.7×10^−6^). ***E***, Distribution of wakeful small and large events over running speed (ns., Kolmogorov-Simonov test).

To estimate the number of APs underlying small and large calcium transients, we generated a look-up table for the number of APs vs. the mean ΔF/F value in the 3-s analysis window from the calcium transients detected in our ground-truth dataset (Fig. 3B, bottom). This relationship was approximately linear for small (<40%) mean ΔF/F values and tapered off at higher values. Note that this tapering-off corresponds to a supralinear increase of ΔF/F values with the number of APs, possibly reflecting additional calcium influx caused by regenerative dendritic events associated with AP bursts (Grienberger et al., 2014; Raus Balind et al., 2019). The variability of the estimated number of APs increased at high mean ΔF/F values, presumably indicating variations of the temporal profile of the underlying spike trains. Applying this look-up table to calcium transients measured during wakefulness, we estimate that small events reflect short bursts of APs or trains of up to 20 APs whereas the largest events with >100% mean ΔF/F presumably were caused by >30 APs within the 3-s window (Fig. 3B, top). As a limitation to this approach, it must be kept in mind that AP patterns, i.e., bursting vs. continuous spiking, are not necessarily preserved between anesthetized and awake states.

The average shape and amplitude of anesthetized calcium transients resembled the small awake events, whereas the awake large events exhibited higher amplitudes and prolonged durations (Fig. 3C). The frequency of small events was comparable during anesthesia and resting wakefulness (p < 0.001, two-sided Wilcoxon rank sum test; Fig. 3D). Behavioral context (resting vs. running) clearly affected the abundance of calcium transients, with significantly increased event frequencies during locomotion compared to resting for both small and large events (Fig. 3D). Large events almost exclusively occurred during running. The distributions of small and large events across running speed were not different (Fig. 3E; p = 0.5, Kolmogorov-Smirnov test), indicating that large events did not simply occur at the highest speeds but rather depended on the behavioral state of the animal.

In summary, we observed especially large-amplitude calcium transients with prolonged duration during wakefulness. Their abundance was particularly high during running behavior, consistent with previous observations that overall spike rates in CA3 pyramidal neurons increase during running (Mizuseki et al., 2012; Oliva et al., 2016).

### Stability and variability of neuronal activity and co-activity in CA3 across days

To assess how stable or variable the activity of CA3 pyramidal neurons is over days, we analyzed calcium transients measured repeatedly in the same neuronal populations over three consecutive days in both anesthetized and awake state (Fig. 4A, B). For each neuron, we calculated the mean ΔF/F peak amplitude, the average inter-event-interval (IEI) time and the average full width half maximum (FWHM) for all calcium transients per day (Fig. 4C, Extended Data Fig. 4-1A, B). We quantified the stability of these features by correlating values recorded during one imaging day with values for the same neurons from the subsequent day (Lütcke et al., 2013) (Fig. 4D). While the ΔF/F amplitude for the same neurons was relatively stable across days (Pearson correlation coefficient ρ = 0.50, 0.34 and 0.54 for anesthetized, resting wakefulness and running conditions; all p < 6×10-7), lower correlation values were found for the features FWHM (ρ = 0.20, 0.09 and 0.15, with p = 0.01, 0.23, 0.02) and IEI (ρ = 0.16, 0.04 and 0.17, with p = 0.09, 0.65, 0.04). Motivated by the observation of relatively stable ΔF/F amplitudes, we specifically addressed the question how the distribution of large events (as defined in Fig. 3) changed over days across the population. Within the subset of neurons (47 neurons out of 182), in which we observed large events on at least one day (33.7% of events in total), about one third of them (33.4%; chance level: 18.2%) displayed large events on at least two days, and a considerable fraction (14.9%; chance level: 0.85%) displayed large events on all 3 consecutive measurement days (Fig. 4E). These above-chance incidences indicate that a subset of neurons exists that is particularly prone to generate large events consistently over days.

**Figure 4.**
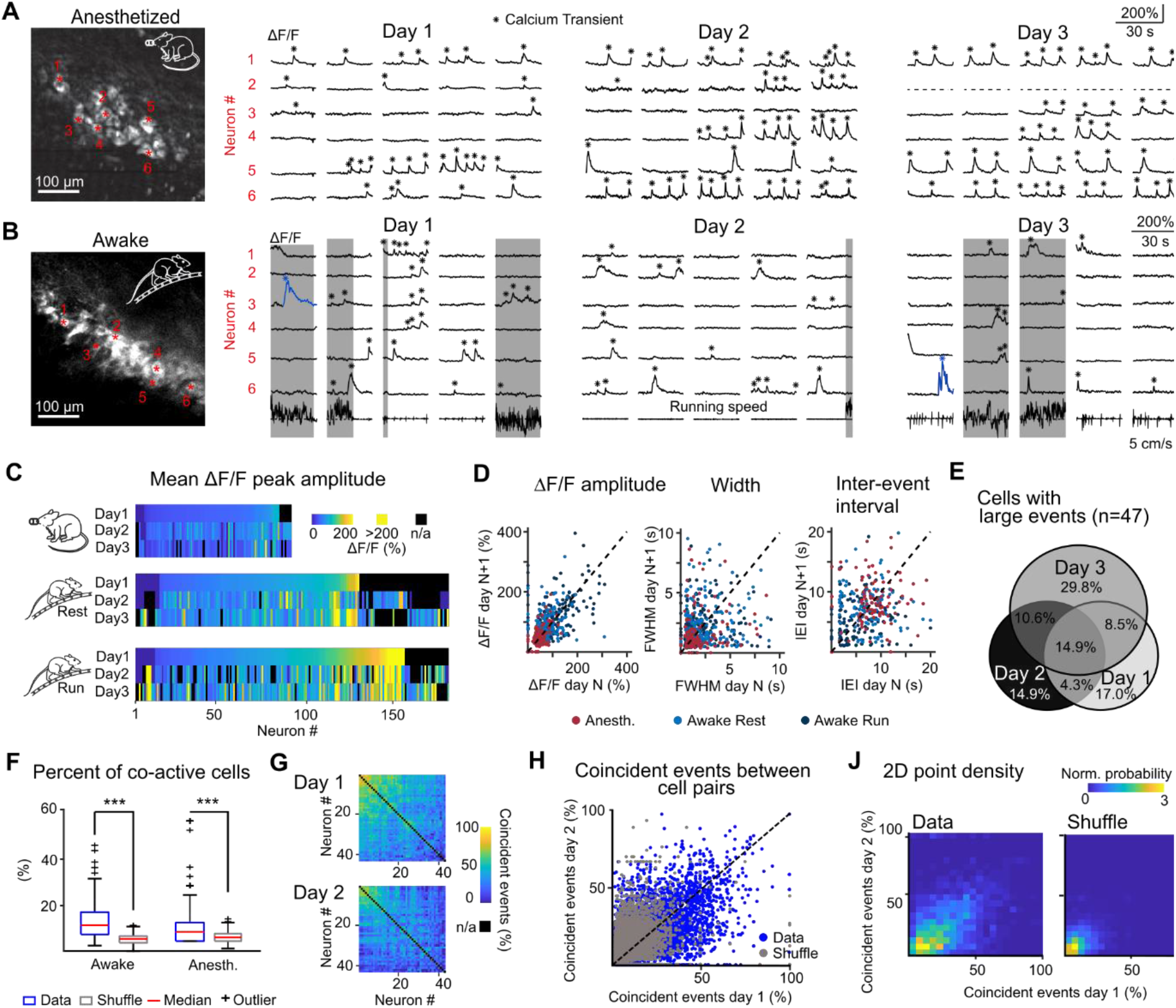
Longitudinal functional imaging of CA3 population activity over consecutive recording days. ***A***, Example fields of view and calcium traces of selected neurons over three consecutive imaging days during anesthesia and ***B***, wakefulness. Large calcium events are labelled blue. Periods of locomotion are highlighted in grey. ***C***, Neuron-wise analysis of mean ΔF/F transient amplitude, three consecutive days in the anesthetized, awake resting and awake running condition. Neurons that were recorded on all three days were sorted according to their properties on the first imaging day (n_anesth._= 91 cells, n_awake_= 182 cells, 6 mice). ***D***, Comparison of mean ΔF/F transient amplitude, mean FWHM and mean inter-event interval per neuron over consecutive imaging days (anesthetized: ρ_Amplitude_=0.50, ρ_FWHM_=0.20, ρ_IEI_=0.16, awake resting: ρ_Amplitude_=0.34, ρ_FWHM_=0.09, ρ_IEI_= 0.04, awake running: ρ_Amplitude_=0.54, ρ_FWHM_=0.15, ρ_IEI_=0.17). ***E***, Percentage of cells firing large events for each day and over consecutive days (n_day1_= 21, n_day2_=21, n_day3_=30, n_day1+day2_ = 9, n_day2+day3_ = 12, n_day1+day3_ =11, n_day1+day2+day3_=7). ***F***, Percentage of neurons per FOV that show coincident calcium transients within a 3 s time window around an event (p < 1×10^−20^ for all event types; two-sided Wilcoxon signed-rank test; ***p<0.001) ***G***, Percentage of coincident events between all neuron pairs of one example FOV over two days. Hierarchical link clustering was performed on the coactivity matrix of day 1 and the resulting order of neurons was maintained for day 2. ***H***, Comparison of percentage of coincident events per neuron pair over consecutive imaging days. For this plot all data from all FOVs was pooled (data: ρ = 0.42, shuffled transient peak times: ρ = 0.29). ***J***, Normalized 2D density plot based on (***H***). <0.001, one-way ANOVA (small events: p_anesth./run_ = 1.7×10^−4^, p_rest/run_ = 4.5×10^−16^; large events: p_rest/run_ = 6.7×10^−6^).

Finally, we investigated CA3 neuronal activity on the population level. To assess synchrony of activity we calculated the percentage of co-active neurons per calcium transient per FOV, with co-activity defined as co-occurrence of calcium transients in a time window ranging from 1 s before to 2 s after an event (Methods). The percentage of co-active neurons was significantly higher than expected from chance level for both anesthetized and awake condition (Fig. 4F; 6.8 ± 5.7% and 11.7 ± 6.9%, respectively; mean ± s.d.; compared to shuffled transient peak times; p < 0.001, Wilcoxon signed-rank test). Additionally, we investigated the stability of co-active neuron pairs by comparing the percentage of coincident calcium transients over two consecutive awake imaging days (Fig. 4G). Percentages ranged from 0 to 91.7% (11.7 ± 2.9 %, mean ± s.d.) and remained relatively stable across two imaging days (data: ρ = 0.42, shuffled transient peak times: ρ=0.29) (Fig. 4H, J), hinting towards functional coupling of neuronal subpopulations in CA3 that is maintained across multiple days.

## DISCUSSION

Our study contributes to the emerging field of in vivo calcium imaging of CA3 pyramidal neurons by establishing longitudinal imaging across days, comparing different behavioral states, and providing calibration in terms of spike patterns underlying the observed calcium transients. We found state-dependent neuronal responses with salient prolonged high-amplitude calcium transients in awake mice during locomotion. On the population level, we observed that during wakefulness individual calcium transients are embedded in surrounding network activity, with coactive neuron pairs maintaining their mutual coactivity over days.

Our juxtacellular recordings during anesthesia and the deconvolved calcium transients from awake imaging sessions indicate low mean firing rates but prominent burst events in CA3 pyramidal neurons, in line with previous studies low mean firing rates but prominent burst events (Henze et al., 2002; Frerking et al., 2005; Wittner and Miles, 2007; Mizuseki et al., 2012; Kowalski et al., 2016; Oliva et al., 2016 Ding et al., 2020). Compared to DG granule cells (Pilz et al., 2016) a much higher fraction of CA3 pyramidal neurons displayed clear calcium transients (>90% for all conditions in CA3; for comparison: <10% during anesthesia and around 50% during wakefulness in DG).

The mean frequency of calcium transients across the entire population was 6- to 20-fold higher in CA3 than in DG, especially during anesthesia. Consistent with the high burstiness of CA3 pyramidal neurons, the vast majority of recorded calcium transients reflected AP bursts rather than individual APs. The bimodal ISI distribution that we observed during anesthesia (Extended Fig. 2-1C) closely resembles previous results during light anesthesia (Kowalski et al., 2016) as well as during sleep and awake behavior (Frerking et al., 2005; Mizuseki et al., 2012). Our data are consistent with state-dependent modulation of AP patterns, with temporally more dispersed APs and reduced burst propensity during locomotion (Mizuseki et al., 2012). However, it is not straightforward to relate the changes in calcium transient frequency that we observed to changes in AP patterns. Moreover, recent in vivo whole-cell recordings found that in most CA3 pyramidal neurons theta oscillations were associated with membrane potential hyperpolarization (Malezieux et al., 2020), which could imply decreased average firing rates during running, contrary to what has been reported previously (Mizuseki et al., 2012; Oliva et al., 2016). In the new study, however, theta periods included both resting and running periods and modulatory effects were quite heterogeneous across the CA3 population. Further investigations will be needed to clarify state-dependent modulation of membrane potential dynamics, AP patterns, and cellular calcium signals in CA3.

We estimate that the especially large and prolonged calcium events that we observed were caused by >30 APs over 3 s, indicating that a subset of CA3 pyramidal neurons can sustain firing rates of 10 Hz or higher during running. This spiking level is not too dissimilar from in-field firing rates observed in identified CA3 place cells (Mizuseki et al., 2012; Ding et al., 2020). As our experiments were conducted in the dark without salient spatial cues, we can only speculate that these events may relate to place cell or time cell properties. Rather than representing regular spiking, we interpret the large locomotion-related events as presumably reflecting a mixture of regular spikes and bursts at shortened inter-burst interval compared to resting conditions (note the ’bumpy’ spike rates in the examples in Fig. 3A; see also Epsztein et al., 2011). Further investigations are required in the future to resolve the electrophysiological basis of these special large events during awake running and their relationship to spatial navigation.

Our juxtacellular recordings indicate that supralinear calcium influx occurs with increasing AP numbers, suggesting additional sources that contribute to the mean ΔF/F values of large events (see Fig. 3D). Additional calcium influx may have been caused by dendritic calcium spikes associated with complex spike bursts (Grienberger et al., 2014; Raus Balind et al., 2019), localized dendritic NMDA spikes (Brandalise et al., 2016), or dendritic plateau potentials induced by supralinear integration of synaptic inputs (Takahashi and Magee, 2009). Plateau potentials and the associated complex spike bursts have been found to precede place field formation in CA1 neurons and may generally mediate behaviorally relevant plasticity in hippocampal pyramidal neurons (Bittner et al., 2017, 2015; Diamantaki et al., 2018).

The recurrent auto-associative nature of the CA3 network is suitable to support the formation of functional neuronal ensembles (Hopfield, 1982; Nakazawa et al., 2002; Guzman et al., 2016). In our experiments, neurons were more frequently co-active during wakefulness compared to anesthesia (Fig. 4G), hinting towards the recruitment of CA3 subpopulations during specific behavioral states or in particular sensory environments. In addition, we found that these co-active ensembles were relatively stable over consecutive days. Previous calcium imaging studies reported unstable space representations on the days-scale in place cells of CA1 and CA3 (Rubin et al., 2015; Hainmueller and Bartos, 2020), although some experiments indicate that representations can be stabilized (Kentros et al., 2004; Mankin et al., 2012; Julian et al., 2018). Despite unstable functional representations in single pyramidal neurons, neurons may maintain a stable affiliation to the same engram (Kinsky et al., 2018; Gonzalez et al., 2019) and spatial information could be stably encoded by whole-network activity patterns, based on pairwise co-activity (Stefanini et al., 2020). Across-day stability of a distributed engram, but variable activation patterns of the pyramidal neurons involved, may allow for flexible functional outputs of a hippocampal subpopulations over time (Goode et al., 2020). In our experiments, a subpopulation of CA3 pyramidal neurons displayed large calcium events consistently across days and showed stable co-activity with other neurons of the same FOV. This result indicates at least some stability in the CA3 neuronal ensemble recruitment processes. Large calcium events associated with complex spike bursting might lead to plasticity in the recurrently connected CA3 network and could support the formation of functional engrams (Raus Balind et al., 2019). The emergence of co-active CA3 ensembles and their relevance for hippocampus-dependent behaviors warrant further investigations using longitudinal calcium imaging.

Because of the fundamental importance of the CA3 subfield in the cortico-hippocampal circuitry, we expect a surge of future in vivo CA3 imaging studies that will be facilitated by recent methodological advances. First, even though two-photon imaging in DG and CA3 has been achieved with GCaMP indicators (Pilz et al., 2016; Hainmueller and Bartos, 2018), red-shifted calcium indicators may still be beneficial (Pilz et al., 2016; Kondo et al., 2017; Shemetov et al., 2020). Second, pushing excitation wavelengths further into the near-infrared wavelength is now possible with three-photon microscopy (Ouzounov et al., 2017), with entire new opportunities for non-invasive hippocampus imaging through the neocortex (Ouzounov et al., 2017; Weisenburger et al., 2019). Finally, the combination of multi-photon imaging with optogenetic manipulation of specific neuronal ensembles, as recently demonstrated in the CA1 region (Robinson et al., 2020), will open new avenues for all-optical interrogation of hippocampal neuronal ensemble dynamics.

## AUTHOR CONTRIBUTION

S.C. and F.H. designed research; G.S. and S.C. performed research; G.S., P.R. and A.A. analyzed data; G.S., P.R. and F.H. wrote the paper

## ACKNOWLEDGEMENT

We thank Lazar Sumanovski for technical assistance and Philipp Bethge, Christopher Lewis and Xiaomin Zhang for comments on the manuscript.

## METHODS

All experimental procedures were conducted in accordance with the ethical principles and guidelines for animal experiments of the Veterinary Office of Switzerland and were approved by the Cantonal Veterinary Office in Zurich.

### Animals and R-CaMP1.07 labelling

For the experiments, male and female mice with a Tg(Grik4-cre)G32-4Stl background were used (n = 6). These mice show a dense expression of Cre-recombinase restricted to CA3 hippocampal pyramidal neurons (MGI:2387441) (Nakazawa et al., 2002). Specific expression of the red fluorescent calcium indicator R-CaMP1.07 (Ohkura et al., 2012) in CA3 pyramidal neurons was achieved by stereotaxic injection of AAV1-EFα1-DIO-R-CaMP1.07 in 6- to 9-week-old adult mice (coordinates: AP -2, ML +1.8, DV -2.2; in mm from bregma; 300 nl with a virus titer of approximately 1×107 vg/nl).

### Hippocampal window implantation

Chronic access for CA3 imaging was obtained by the implantation of a hippocampal window (Pilz et al., 2016). One week after the virus injection we performed a 3-mm diameter craniotomy centered at the injection site and implanted a stainless-steel cannula with a front glass window. After removing the bone, we gently aspirated the underlying cortical tissue until the corpus callosum fibers became visible. A stainless steel cannula (Ø 3 mm, 1.5 mm length) covered by a glass coverslip (Ø 3 mm, 0.17 mm thickness) was inserted into the cavity and secured in place using dental acrylic cement (Ivoclar vivadent) (Fig. 1A, B). Additionally, an aluminium post for head fixation during imaging was attached to the skull. After a recovery period, mice were handled by the experimenter, habituated to head fixation, and accustomed to run on a ladder wheel (Ø 20 cm) with regularly spaced rungs (1-cm spacing) during head fixation. Approximately two weeks after the surgery, neuronal population activity was imaged under isoflurane anesthesia (1-2% in oxygen) on three to five consecutive days. The same neuronal populations that were imaged in the anesthetized condition were repeatedly imaged in awake animals for 5-10 days, of which at least 3 days were consecutive (Fig. 1C).

### Two-photon calcium imaging

We used a custom-built two-photon microscope based on the Sutter Movable Objective Microscope (MOM) type, equipped with a water immersion 16x objective (CFI LWD 16X/0.80; Nikon), a Pockels cell (model 350/80 with controller model 302RM, Conoptics), and galvanometric scan mirrors (model 6210; Cambridge Technology), controlled by HelioScan software (Langer et al., 2013). R-CaMP1.07 was excited by ∼230-fs pulses at 80 MHz provided by a ytterbium-doped potassium gadolinium tungstate (Yb:KGW) laser (1040 nm; >2 W average power; model Ybix; Time-Bandwidth Products). Emitted fluorescence was detected by a photomultiplier tube after passing through a 610/75 nm bandpass filter (AHF Analysetechnik). Laser intensities during imaging were 56-78 mW under the objective.

In anesthetized experiments, mice were anesthetized with isoflurane (1-2% in oxygen). Body temperature was monitored continuously with a thermosensor and kept at 37°C with a heating blanket. For awake experiments, the head-fixed mouse was placed on the ladder wheel and was free to run. Running speed and running distance during calcium imaging were recorded at 40 Hz with a rotary encoder (Phidgets, 12V/0.2Kg-cm/230RPM 10:1 DC gear motor with encoder). The activity of R-CaMP1.07-expressing CA3 pyramidal cells was recorded in trials of 30-s duration, with 10-s inter-trial intervals (maximum of 30 trials per day). Recordings were performed in the distal part of CA3, which lays in the proximity of CA2. In all sessions, imaging across a field of view (FOV) of 325 x 325 µm2 was performed at 10 Hz frame rate.

### *In vivo* electrophysiology

Electrophysiological recordings, combined with in vivo calcium imaging, were performed in acute in vivo preparations of Tg(Grik4-cre)G32-4Stl expressing R-CaMP1.07 mice (n = 3; at least two weeks after injection). Mice were anesthetized with isoflurane and the temperature was maintained at 37°C. A stainless steel plate was fixed to the exposed skull using dental acrylic cement. A 4-mm diameter craniotomy was performed, centered above the virus injection locus. The overlying cortex was aspirated until the corpus callosum became visible. A 1%-agarose gel was filled into the cavity to reduce tissue motion. Juxtacellular recordings from R-CaMP1.07-expressing CA3 pyramidal neurons were obtained with glass pipettes (4-6 MΩ pipette resistance) filled with Ringer’s solution. To facilitate visually-guided targeting of individual neurons, the pipette was coated with BSA Alexa-594 (Invitrogen). Action potentials were recorded juxtacellulary in current clamp mode at 10-kHz sampling rate using an Axoclamp 2B amplifier (Axon Instruments, Molecular Devices) and digitized using Clampex 10.2 software. Simultaneously, we performed two-photon calcium imaging of the recorded neuron at 10 Hz frame rate.

### Perfusion and histology

After the last awake imaging session, mice were administered a lethal dose of pentobarbital (Ekonarcon, Streuli) and transcardially perfused with sterile NaCl (0.9%) followed by 4% paraformaldehyde (PFA, 0.1 M phosphate buffer, pH 7.4). 40-µm coronal brain slices were obtained and histological images were acquired with a confocal laser-scanning microscope (Olympus FV1000) using 546-nm laser light for R-CaMP1.07 excitation (Fig. 1B).

### Data analysis

Electrophysiological data were analyzed using routines in IGOR (Wavemetrics). R-CaMP1.07 fluorescence signals were analyzed using custom-written macros in ImageJ (Schindelin et al., 2012) and MATLAB routines (Mathworks). For motion correction of calcium imaging movies, we applied a Hidden Markov Model line-by-line motion correction algorithm (Dombeck et al., 2007). We excluded trials that obviously were insufficiently motion-corrected based on visual inspection. Regions of interest (ROIs) corresponding to individual neurons were manually selected from the mean fluorescence image of a single-trial time series. Background fluorescence was estimated as the bottom 1st percentile fluorescence signal across the entire session and subtracted before calculating the relative percentage fluorescence change from baseline ΔF/F = (F-F0)/F0. Baseline fluorescence F0 was computed as 51st percentile of the fluorescence signal in an 8-s sliding window. ΔF/F traces were smoothed with a 5-point 1st-order Savitsky-Golay filter. The simultaneously recorded electrical and fluorescence signals were aligned at the start of recording. We determined the peak amplitude of the recorded calcium transient and counted the number of underlying APs (burst events within APs maximally spread over a 200-ms period were included in this analysis). For averaging, calcium transients were aligned to the first AP of a given event. Detectable calcium transients were defined as fluorescence signals that deviated from baseline by >3 s.d. for anesthetized imaging and >4 s.d. for awake conditions (more stringent criterion for awake conditions because of increased noise levels and possible motion artefacts during wakefulness). For every threshold crossing event we determined the calcium transient peak as the first maxima (using the MATLAB function findpeaks with a minimal peak prominence of 20% ΔF/F and a minimal distance between peaks of 1.5 s).

We excised 3-s segments around detected calcium transient events (−1 s to +2 s relative to the peak) and deconvolved the calcium transient using a supervised algorithm based on neural networks (Rupprecht et al., 2020). The algorithm was trained on the simultaneous recordings of action potentials and calcium transients (n = 4 neurons from 3 mice, a total of 33 minutes of recording and 5025 APs), with the ground truth data re-sampled to the target frame rate (10 Hz). Using additive Poisson noise, the noise level of the ground truth was adjusted to match the noise level of each neuron of the population imaging data. The ground truth recorded during anesthesia did not cover calcium transient amplitudes and shapes representative of the large and prolonged calcium transients observed during wakefulness. To estimate the number of action potentials during these calcium transients in Fig. 3, we therefore used a model-free look-up table based on the integral of the spike rate predictions in the excised 3-s calcium transient segment. For the awake recordings, we defined ‘large’ calcium transients as those that displayed mean F/F values in the 3-s time window larger than the 95th percentile of the distribution of mean F/F values for all transients recorded during anesthesia.

For analysis of neuronal population activity, we determined the level of co-activity of neuron pairs (e.g., neuron A and neuron B) by analyzing for each detected calcium transient in neuron A whether a calcium transient also occurred in neuron B in a surrounding 3-s window (1 s before to 2 s after the transient peak in neuron A). Co-activity of neuron pairs, defined as the probability that two neurons of the same FOV showed coincident calcium transients in this 3-s window, was determined for the awake and the anesthetized condition and compared to the co-activity of neuron pairs based on shuffled time points of calcium transient peaks. Experimental data and shuffled data were pooled from all FOVs and compared using Wilcoxon signed rank tests.

### Run speed analysis

We down-sampled running speed to the 10-Hz imaging frame rate and defined periods with >0.5 cm/s speed as ‘run’ periods and periods with lower speeds as ‘rest’ periods. The numbers of small and large calcium transients per minute during wakeful resting or locomotion were determined and distributions were compared using one-way ANOVA. The distributions of run speed of wakeful small and large events were compared using a Kolmogorov-Smirnov test.

## EXTENDED DATA

**Extended Data Figure 2-1**

**Figure 2-1.**
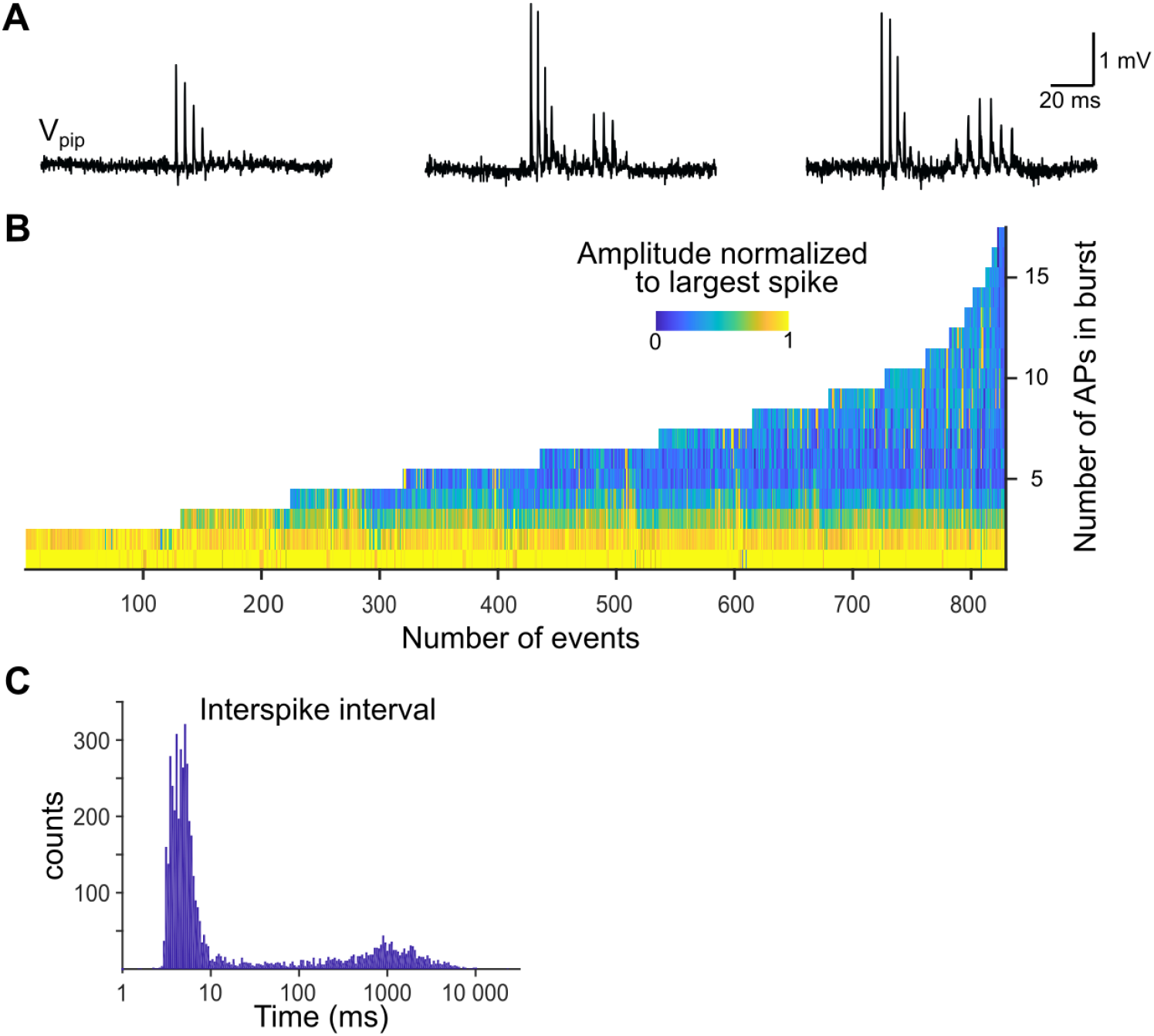
Heterogeneity of burst events in CA3 pyramidal neurons from cell-attached ground-truth recordings. **A**, Electrophysiological recordings of three example bursts. Example two and three show intermittent bursts. **B**, Overview of all recorded bursts with up to 17 APs. Amplitudes were normalized to the maximum AP amplitude within the burst. **C**, Distribution of inter spike interval times pooled from all recordings (4 neurons, 3 mice).

**Extended Data Figure 4-1**

**Figure 4-1.**
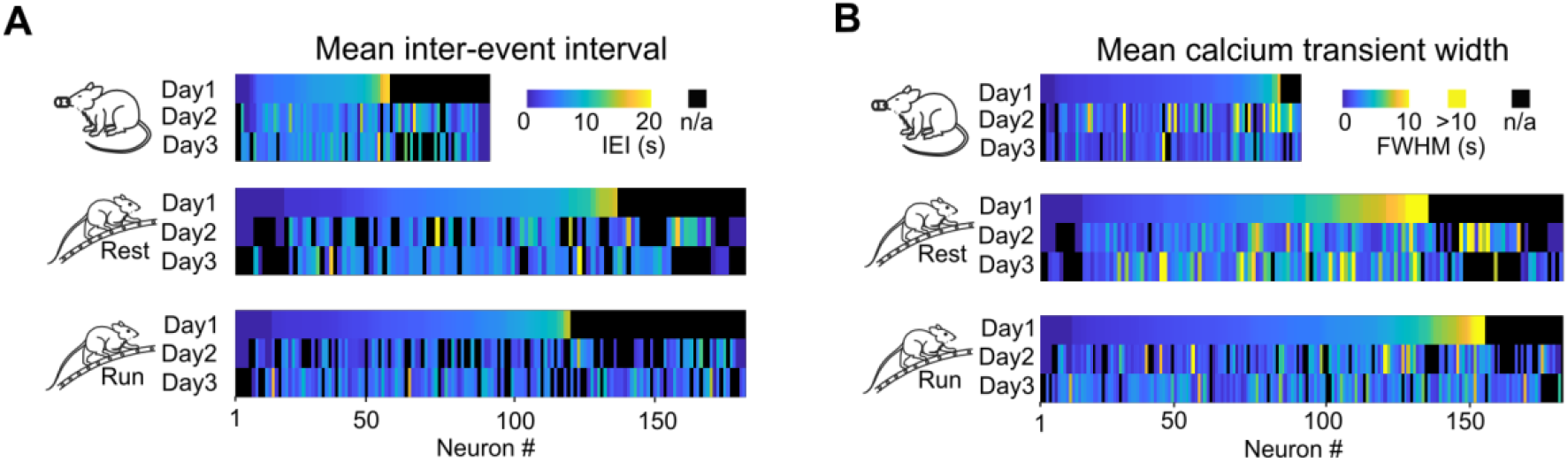
Properties of calcium transients of CA3 neurons over consecutive imaging days. Neuron-wise analysis of ***A***, mean inter-event interval and ***B***, mean transient FWHM on three consecutive days in the anesthetized, awake resting and awake running condition. Neurons that were recorded on all three days were sorted according to their properties on the first imaging day (n_anesth._= 91 cells, n_awake_= 182 cells, 6 mice).

